# Naa10p promotes metastasis by stabilizing matrix metalloproteinase-2 protein in human osteosarcomas

**DOI:** 10.1101/243592

**Authors:** Ming-Hsien Chien, Wei-Jiunn Lee, Yi-Chieh Yang, Peng Tan, Ke-Fan Pan, Hsiao-Chi Tsai, Chun-Hua Hsu, Michael Hsiao, Kuo-Tai Hua

**Affiliations:** Graduate Institute of Clinical Medicine, College of Medicine, Taipei Medical University, Taipei, Taiwan; Department of Medical Education and Research, Wan Fang Hospital, Taipei Medical University, Taipei, Taiwan; Department of Urology, School of Medicine, Taipei Medical University, Taipei, Taiwan; The Genomics Research Center, Academia Sinica; Taipei, Taiwan; Graduate Institute of Toxicology, College of Medicine, National Taiwan University, Taipei, Taiwan; Graduate Institute of Basic Medical Science, China Medical University, Taichung, Taiwan; Department of Agricultural Chemistry, National Taiwan University, Taipei, Taiwan; Department of Biochemistry, College of Medicine, Kaohsiung Medical University, Kaohsiung, Taiwan

**Keywords:** Naa10p, MMP-2, protein stability, osteosarcoma, metastasis

## Abstract

N-α-Acetyltransferase 10 protein (Naa10p) mediates N-terminal acetylation of nascent proteins. Oncogenic or tumor suppressive roles of Naa10p were reported in cancers. Here, we report an oncogenic role of Naa10p in promoting metastasis of osteosarcomas. Higher *NAA10* transcripts were observed in metastatic osteosarcoma tissues compared to non-metastatic tissues and were also correlated with a worse prognosis of patients. Knockdown and overexpression of Naa10p in osteosarcoma cells respectively led to decreased and increased cell migratory/invasive abilities. Re-expression of Naa10p, but not an enzymatically inactive mutant, relieved suppression of the invasive ability *in vitro* and metastasis *in vivo* imposed by Naa10p-knockdown. According to protease array screening, we identified that matrix metalloproteinase (MMP)-2 was responsible for the Naa10p-induced invasive phenotype. Naa10p was directly associated with MMP-2 protein through its acetyltransferase domain and maintained MMP-2 protein stability via NatA complex activity. MMP-2 expression levels were also significantly correlated with Naa10p levels in osteosarcoma tissues. These results reveal a novel function of Naa10p in the regulation of cell invasiveness by preventing MMP-2 protein degradation that is crucial during osteosarcoma metastasis.

## Introduction

Osteosarcomas are the most common primary malignant tumor of the bone in children and adolescents.[1] Although clinical management of osteosarcomas, including surgery and chemotherapy, has significantly improved long-term survival over the past few decades, outcomes for those patients with metastatic or recurrent osteosarcoma remain dismally poor and, therefore, novel therapeutic strategies are urgently required.[2]

Matrix metalloproteinases (MMPs) play important roles in the progression of several types of cancer by increasing tumor growth, migration, invasion, and metastasis and are associated with poor disease prognosis.[3] Tumor growth of osteosarcomas is accompanied by both enhanced local bone destruction and bone formation, two processes that are dependent on proteolytic enzymes. Indeed, many MMPs were found to be expressed in primary and metastatic osteosarcoma tissues.[4] MMP-2, -9, and -14 as well as tissue inhibitor of matrix metalloproteinase (TIMP)-1 were shown to be associated with a poor prognosis of osteosarcomas.[5, 6] Intense immunostaining for MMP-2 and -9 was also detected in metastatic lesions from osteosarcomas.[4, 7] However, MMP-2 was recently considered to be a more-important MMP involved in osteosarcoma progression. A shift from MMP-9 to predominant MMP-2 activity was observed in zymographic screening of cryo-preserved osteosarcoma biopsies and was associated with a poor response to chemotherapy.[8]

N-α-Acetyltransferase 10 protein (Naa10p) is the catalytic subunit of a heterodimeric complex, NatA, which co-translationally acetylates N termini of small side-chain amino acids such as Ser, Ala, Thr, Gly, Val, and Cys, after the initiator methionine has been cleaved from nascent polypeptides.[9, 10] Despite being debated, Naa10p was also found to acetylate proteins at internal lysine residues of diverse targets.[11] Through N-α or N-ε acetylation, Naa10p regulates protein stability[12], protein activity[13], and protein-protein interactions[14, 15] in a variety of target proteins. Non-acetylation-dependent functions of Naa10p were also documented in regulating cell behaviors.[16, 17] Thus, Naa10p seems to be a diverse protein with different capacities for executing its effects on target proteins. This diversity also reflects the identified roles of Naa10p in cancer cells. Dependent on the cell context, Naa10p is involved in regulating cell proliferation, apoptosis, autophagy, metastasis, and chemosensitivity of different cancer cells.[9] Naa10p overexpression was reported in breast[18], colorectal[19], liver[20], and lung cancers[21], while low expression in non-small cell lung cancer (NSCLC) was also documented[12]. Furthermore, loss of heterozygosity at Naa10p gene loci was also found in tumors including lung, breast, pancreatic, and ovarian cancers.[12] Considering the diverse targets Naa10p may regulate in different types of cancer cells or in different stages during cancer development, identifying cancer-type specific targets may help to understand the role of Naa10p in a particular cancer type.

Recently, Naa10p was found to participate in controlling osteoblast differentiation and bone formation as a feedback regulator of Runx2.[22] Naa10p transgenic mice showed delayed calvarial bone development, while Naa10p-knockout mice showed facilitated calvarial bone development. Naa10p-knockdown also augmented the healing of rat calvarial defects. These results highlight a negative role of Naa10p in osteoblast differentiation and bone development. However, whether Naa10p is involved in the formation or progression of osteosarcomas is completely unknown. Herein, we showed that Naa10p is functionally involved in the invasion and metastasis of osteosarcomas. Naa10p is highly expressed in late-stage osteosarcomas and was correlated with shorter survival in patients. Mechanistic studies revealed that Naa10p directly interacts with MMP-2. Naa10p increases MMP-2 protein stability in an N-α-acetylation-dependent manner and thus increases the invasive ability of osteosarcoma cells. Our findings highlight the potential of targeting Naa10p-MMP2 regulation in therapeutic applications against osteosarcomas.

## Results

### Naa10p expression correlated with poor prognosis of osteosarcoma patients and invasiveness of osteosarcoma cell lines

Naa10p was recently reported to be expressed by osteoblasts which are located around new bone surfaces.[22] To examine the role of Naa10p in osteosarcomas, which arise from a mesenchymal origin and exhibit osteoblastic differentiation, we analyzed Naa10p expression in a tissue microarray composed of osteosarcomas at different clinical stages. Our results showed that Naa10p was barely expressed by normal cartilage; on the contrary, Naa10p expression was enriched in osteosarcoma tissues (Figure 1a). Furthermore, Naa10p seemed to be expressed at higher levels in advanced stages compared to early stages of osteosarcomas. To further examine the clinical importance of Naa10p expression in osteosarcomas, we analyzed correlations of Naa10p messenger (m)RNA expressions with metastasis and survival probabilities from a publicly available osteosarcoma cohort, GSE21257. Naa10p mRNA levels were significantly higher in specimens collected from osteosarcoma patients with metastasis at the time of diagnosis than from osteosarcoma patients without metastasis at the time of diagnosis (Figure 1b). Moreover, osteosarcoma patients with above average Naa10p expression levels showed significantly shorter overall survival periods (*p* = 0.032, Figure 1c). These data suggest that Naa10p may play an important role in osteosarcoma progression. We next asked whether the increase in Naa10p expression during osteosarcoma progression was correlated with advanced cell behaviors. A screening of Naa10p expression in six osteosarcoma cell lines showed different levels of Naa10p expression (Figure 1d). These cell lines also showed different degrees of cell migratory and invasive abilities (Figure 1e, f). Interestingly, Naa10p expression was higher in cells with high cell mobility (HOS, MNNG/HOS, 143B, and U2OS cells) and lower in cells with low cell mobility (MG-63 and Saoa-2 cells). Collectively, these results suggest that Naa10p expression is highly correlated with the aggressiveness of osteosarcomas as examined in both cell models and clinical patients.

**Figure 1.**
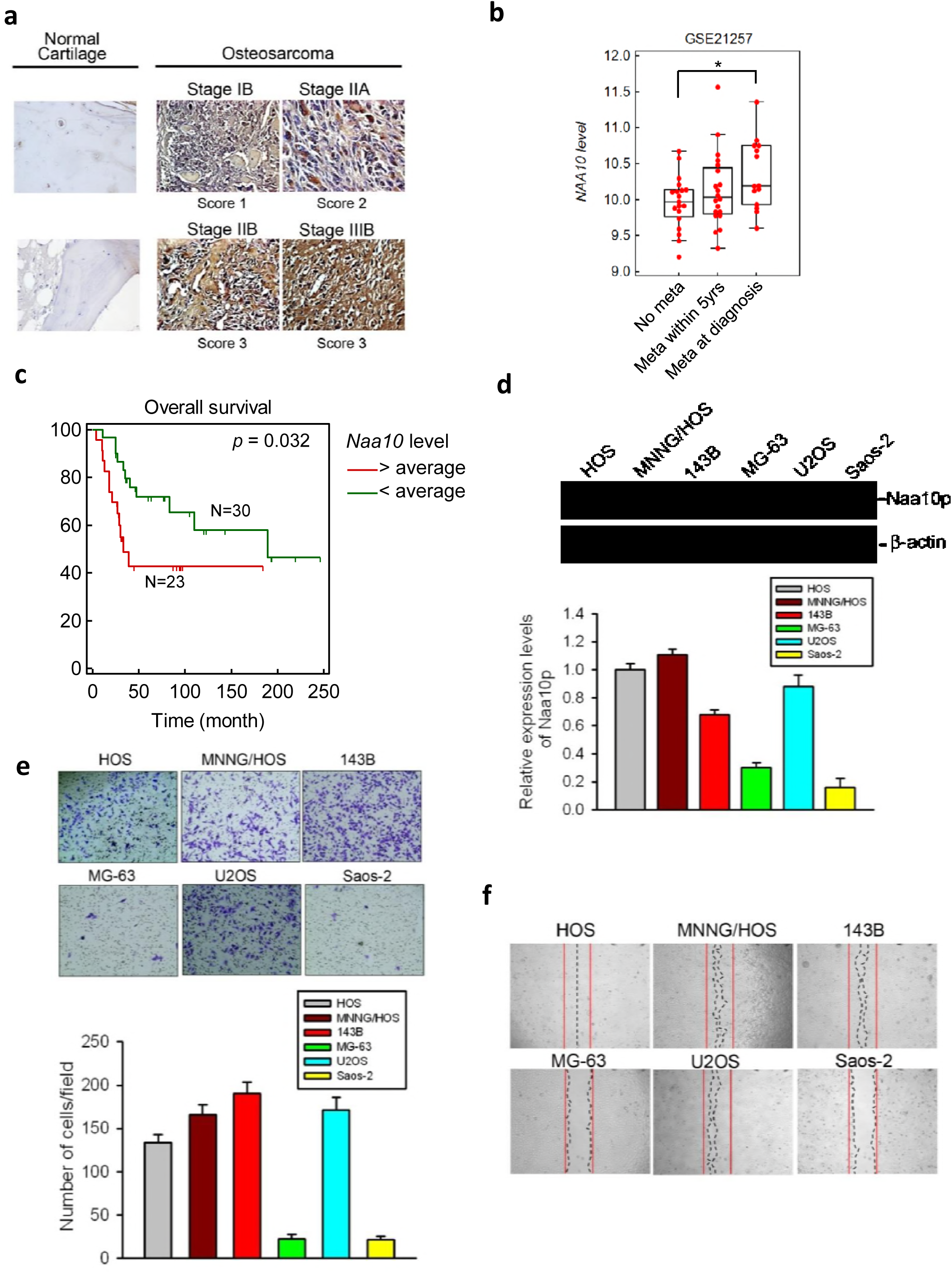
Naa10p expression correlates with poor prognosis of osteosarcoma patients and invasiveness of osteosarcoma cell lines. (**a**) Representative microphotographs of Naa10p immunohistochemical staining in normal cartilage and osteosarcoma tissues from different clinical stage. (**b**) Plot depicting expression levels of *Naa10p* in osteosarcoma specimens from patients with either no metastasis, metastasis within 5 years, or metastasis at diagnosis. The plot was made using GSE21257. (**c**) Kaplan-Meier plot showing the survival of 53 osteosarcoma patients with either high (> average) or low (<average) expression of *Naa10p*. (**d**) Top: Protein expression of Naa10p in osteosarcoma cell lines. Bottom: Quantitative results of Naa10p protein levels which were normalized to β-actin protein levels. Values are presented as the mean ± SE of three independent experiments. (**e**) Invasive abilities of osteosarcoma cell lines as determined by transwell invasion assays. Top: Representative microphotographs of transwell assays. Bottom: Quantitative results by counting invaded cells in a 100× field. Three random fields were counted per well. (**f**) Representative micrographs of wound-healing migration assays were photographed using a phase-contrast microscope (100×).

### Naa10p promotes migratory/invasive abilities of osteosarcoma cells

We next evaluated the importance of Naa10p in regulating cell migration and invasiveness of osteosarcoma cell lines. Knockdown of Naa10p was performed by two specific short hairpin (sh)RNAs in highly invasive 143B and MNNG/HOS cells (Figure 2a). The invasive abilities of control and Naa10p-knockdown cells were next estimated with a transwell invasion assay. As shown in Figure 2b, significantly fewer invaded cells of both cell lines were seen in Naa10p-knockdown groups than in control groups. Moreover, Naa10p-knockdown also significantly decreased the cell migratory ability of MNNG/HOS cells (Figure 2e, left panel), and these results suggest an essential role for Naa10p in maintaining high cell motility of osteosarcoma cells. In comparison, we further overexpressed V5-tagged Naa10p in poorly invasive MG-63 and Saos-2 cells (Figure 2c) and evaluated their cell invasive abilities. Compared to vector control cells, Naa10p-overexpressing cells showed significant increases in the invasive and migratory abilities (Figure 2d, e, right panel). To confirm the dependence of Naa10p in regulating cell invasion and exclude possible off-target effects of shRNA, we re-expressed Naa10p in Naa10p-depleted MNNG/HOS cells and examined their invasive ability. Naa10p-V5 overexpression significantly relieved the Naa10p-knockdown-induced decrease in the invasive ability (Figure 2f). None of the results from migration and invasion assays was affected by possible differences in cell viability, since no viability changes were observed within 48 h between the control and experimental groups (Figure 2g). Taken together, these *in vitro* assays demonstrated that Naa10p expression is essential and sufficient to maintain the high invasive ability of osteosarcoma cells.

**Figure 2.**
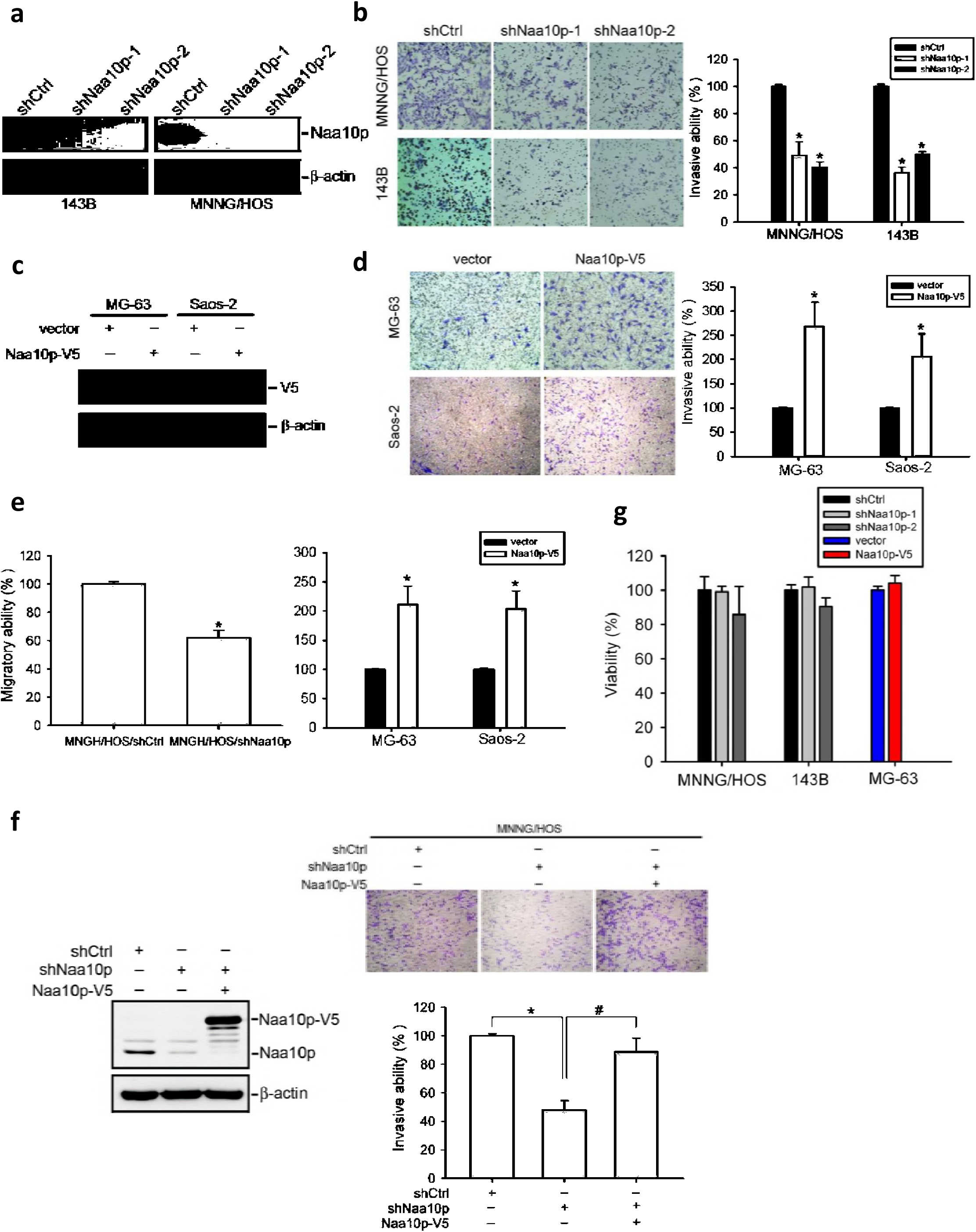
Naa10p promotes the migratory/invasive abilities of osteosarcoma cells. (**a**) Knockdown efficiencies of two Naa10p shRNAs were determined by Western blotting in 143B and MNNG/HOS cells. (**b**) Invasive abilities of Naa10p-knockdown 143B and MNNG/HOS cells were examined by transwell invasion assays. Cells invaded into the opposite site of the transwell were counted, and data are normalized to percent of control. Representative micrographs of invasion assays shown on the left. (**c**) Naa10p was overexpressed in MG-63 and Saos-2 cells as determined by Western blotting. (**d**) Invasive abilities of Naa10p-overexpressing MG-63 and Saos-2 cells were determined by a transwell invasion assay. Cells that had invaded into the opposite site of the transwell were counted, and data are normalized to percent of control. Representative micrographs of invasion assays are shown on the left. (**e**) Transwell migration study of Naa10p-manipulated cells. Cells that had migrated into the opposite site of transwell were counted, and data are normalized to percent of control. (**f**) Transwell invasion assays of MNNG/HOS cells expressing Naa10p with or without co-expressing Naa10p-V5 as indicated. Expressions of endogenous and overexpressed Naa10p were determined by Western blotting on the left. Representative micrographs of invasion assays are shown above the quantitative plot. (**g**) Changes in cell viability of Naa10p-manipulated cells were determined by MTS assays. Data are shown as a percent of the control cells. Values are presented as the mean ± SE of three independent experiments. * *p* < 0.05, compared to control cells. # *p* < 0.05, compared to Naa10p shRNA-infected MNNG/HOS cells.

### Naa10p regulates tumorigenicity and metastatic ability in an animal model

Our clinical analysis and *in vitro* experiments suggested the possible involvement of Naa10p in promoting osteosarcoma metastasis. We next evaluated the *in vivo* effects of Naa10p expression on tumor growth and metastasis. Luciferase-expressing MNNG/HOS/shCtrl and MNNG/HOS/shNaa10p cells were established and implanted orthotopically into the proximal tibia of NOD/SCID mice. Tumor growth and metastasis were monitored through bioluminescence imaging. Control MNNG/HOS cells (MNNG/HOS/shCtrl) orthotopically injected into SCID mice produced larger tumors than did MNNG/HOS/shNaa10p cells injected in the mice after 31 days, as revealed by photon emission detection (Figure 3a). Tumor weights of removed xenografts from the Naa10p-knockdown group were also significantly smaller than those of the control group (Figure 3b). Moreover, we detected Naa10p expression of xenografts harvested from MNNG/HOS/shCtrl-or MNNG/HOS/shNaa10p-injected mice and observed that Naa10p expression was still lower in Naa10-knockdown xenografts compared to control xenografts (Figure 3c). About 90% of distant metastases of osteosarcomas occur in the lungs.[23] *Ex vivo* lung images from MNNG/HOS/shNaa10p-injected mice exhibited a significantly lower intensity compared to those in MNNG/HOS/shCtrl mice (Figure 3d). Furthermore, we intravenously injected MNNG/HOS/shCtrl and MNNG/HOS/shNaa10p cells through the tail vein of NOD/SCID mice and evaluated their survival probability. As shown in Figure 3e, 70% mice from the Naa10p-knockdown group were still alive 40 days after the injection, while only 23% of mice from the control group were still alive. These data support the essential role of Naa10p in sustaining tumor growth and maintaining high metastatic features of osteosarcoma cells.

**Figure 3.**
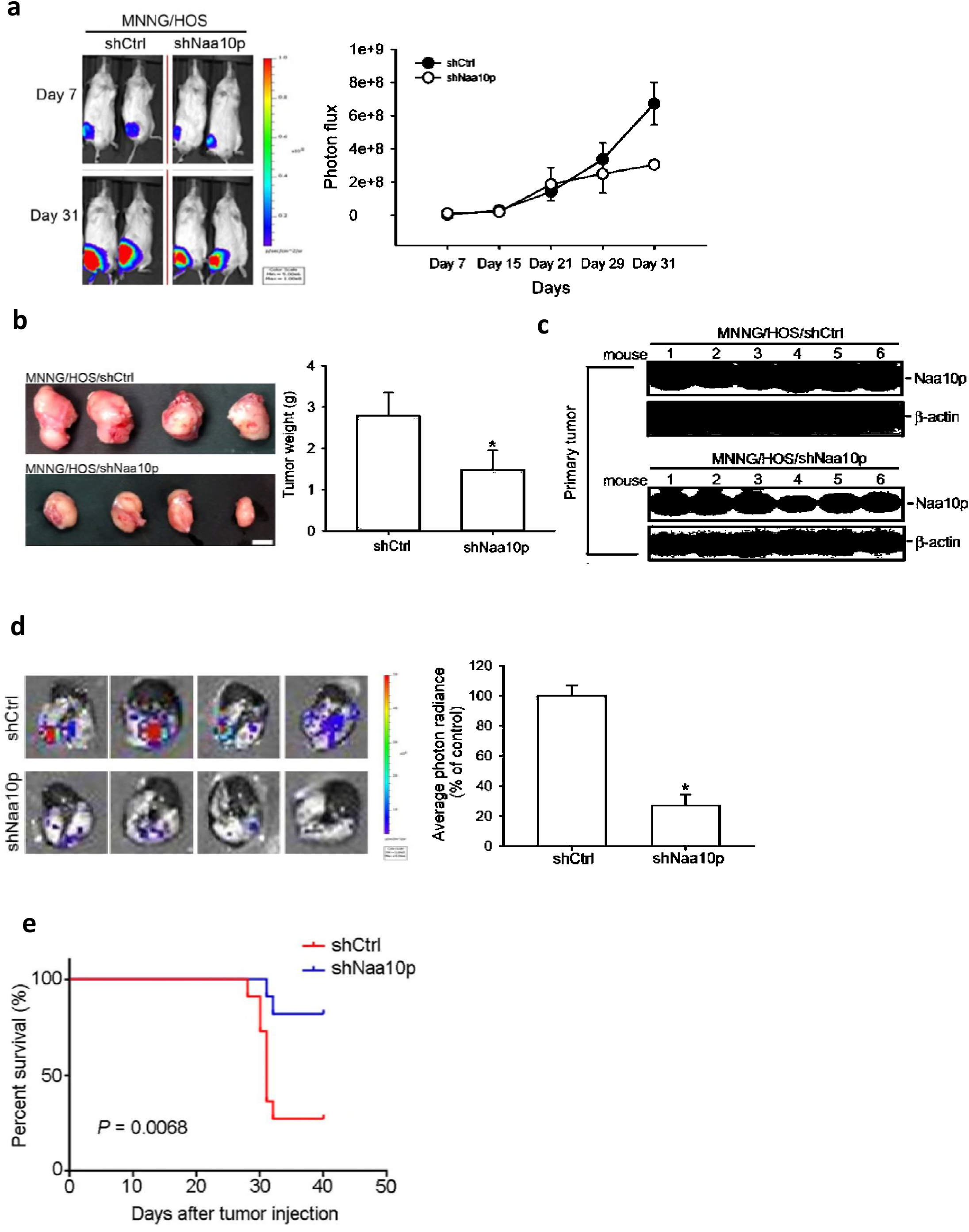
Naa10p regulates tumorigenecity and metastatic ability *in vivo*. (**a**) MNNG/HOS cells (5 × 10^5^) were injected into the proximal tibia of NOD/SCID mice (six mice/group). Representative bioluminescent images of mice from the control and shNaa10p groups taken on days 7 and 31 are shown on the left. Tumor growth rates were estimated by analyzing bioluminescent activities at different time points. Thirty-one days after tumor cell injection, animals were sacrificed, and tumor specimens were removed for further analysis. (**b**) Primary tumors were excised, and representative photos are shown on the left. Tumor weights were measured and are shown on the right. (**c**) Naa10p expressions in excised primary tumors from the control and shNaa10p groups were determined by Western blotting. (**d**) Lungs were isolated and examined from the orthotopic xenograft model. Left: Representative luciferase activity image. Right: Quantification of luciferase activity. (**e**) MNNG/HOS cells (10^6^) expressing shNaa10p or shCtrl were tail vein injected into NSG mice. Survival rates between these two groups (11 mice per group) were monitored and analyzed by a Kaplan-Meier plot. Significance was determined by the log-rank test. * *p* < 0.05. Error bars indicate the SE.

### Naa10p regulates osteosarcoma invasion through stabilizing the MMP-2 protein

Since Naa10p regulated cell invasion of the osteosarcoma cell lines, we performed proteomic screening with a human protease array (ARY021, R&D Systems), which contained 34 different proteases, to investigate the underlying mechanism. Several proteases were decreased in Naa10p-depleted MNNG/HOS cells compared to control cells (Figure 4a). We next validated the results by Western blot analysis of MMP-2, MMP-3, MMP-9 and presenilin-1 in MNNG/HOS/shCtrl and MNNG/HOS/shNaa10p cells and found that only MMP-2 and MMP-9 protein levels decreased after Naa10p depletion (Figure 4b and Supplementary Figure 1a). MMP-2 was reported to express in all osteosarcoma cells from tissue specimens and established cell lines and is known to associate with osteosarcoma metastasis.[24, 25] MMP-2 expression was also increased after Naa10p overexpression in MHHG/HOS cells (Figure 4b). Additionally, we observed that MMP-2 expression in tumor specimens dramatically decreased in the Naa10p-knockdown group compared to the control group (Figure 4c). To test the importance of MMP-2 expression in Naa10p-promoted cell invasion, we rescued MMP-2 in Naa10p-knockdown MHHG/HOS cells by transfection of an MMP-2-expressing plasmid (Figure 4d, upper panel). MMP-2 overexpression alone significantly increased the invasive ability and also rescued the invasion suppression imposed by Naa10p-knockdown in MHHG/HOS cells (Figure 4d), suggesting the dependence of Naa10p-regulated cell invasiveness on MMP-2. Since Naa10p was previously found to regulate the protein stability of target proteins[12, 26], we next tested whether MMP-2 protein stability was affected by Naa10p expression. The MMP-2 protein half-life was estimated in MNNG/HOS/shCtrl and MNNG/HOS/shNaa10p cells by examining protein expression levels after cycloheximide (CHX) treatment. MMP-2 protein quickly diminished after CHX treatment in MNNG/HOS/shNaa10p cells with an estimated half-life of around 1 h. However, the MMP-2 protein half-life was more than 8 h in MNNG/HOS/shCtrl cells (Figure 4e). Furthermore, we also observed that pretreatment with the proteasome inhibitor, MG-132, was able to prevent MMP-2 degradation after Naa10p-knockdown (Figure 4f). Therefore, Naa10p may regulate the MMP-2 protein stability through preventing MMP-2 from proteasome-dependent protein turnover. We next investigated whether MMP-2 expression was correlated with Naa10p levels in human osteosarcoma patients. Immunohistochemical (IHC) staining of osteosarcoma specimens revealed a significant correlation between MMP-2 and Naa10p expressions (Figure 4g).

**Figure 4.**
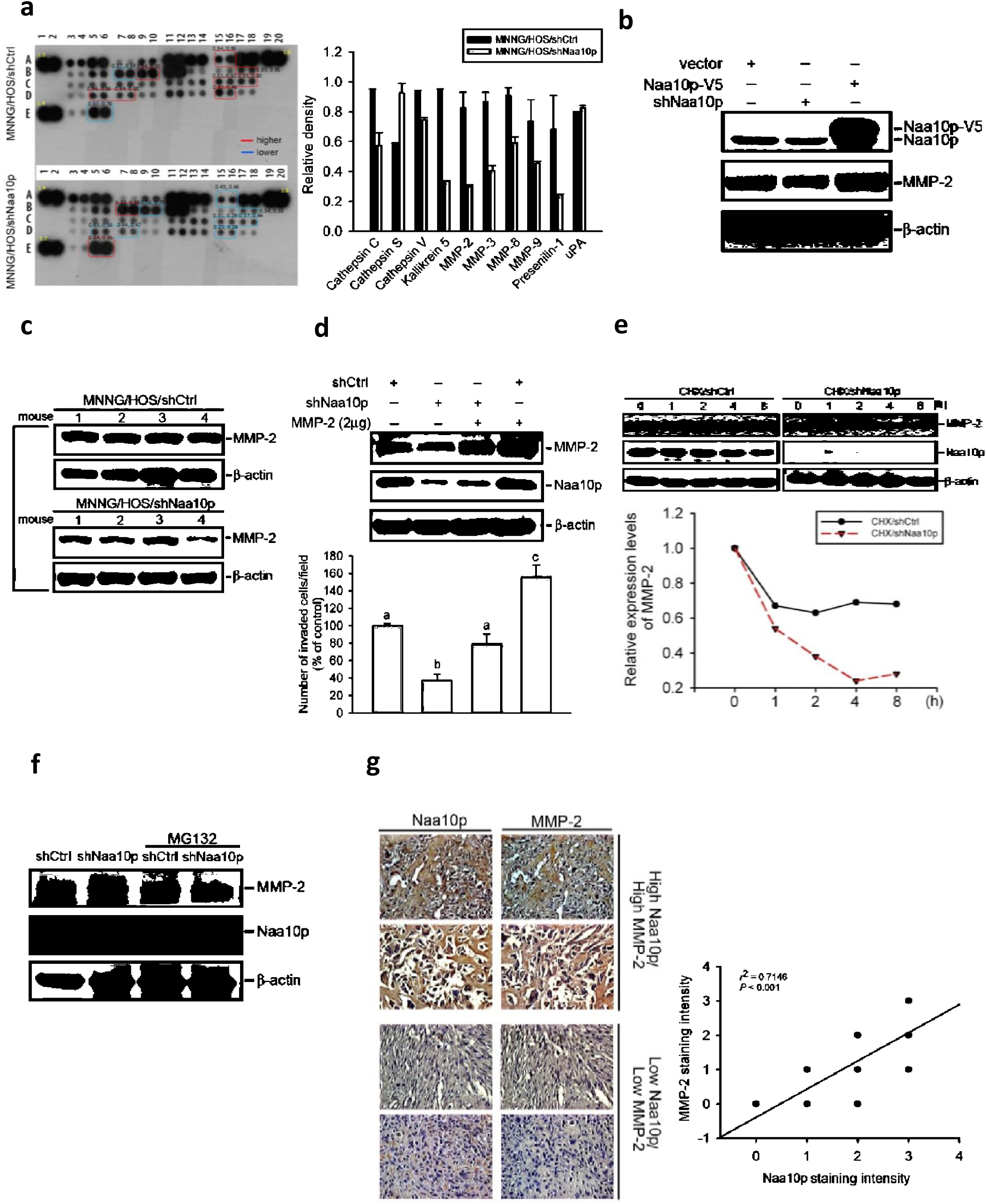
Naa10p regulates osteosarcoma invasion through stabilizing the matrix metalloproteinase (MMP)-2 protein. (**a**) Changes in expressions of proteases in MNNG/HOS/shCtrl and MNNG/HOS/shNaa10p cells. An antibody array (R&D Systems) containing 34 different antibodies against proteases was used to evaluated proteases expression. Left panel showed the representative array blots Right panel showed the quantitative analysis of the protease array Values are presented as the mean ± SE. *n* = 2. (**b**) MNNG/HOS cells expressed shNaa10p with or without co-expression of Naa10p-V5 as indicated. Cell lysates were harvested, and MMP-2, Naa10p, and β-actin levels were determined by Western blotting. (**c**) Protein lysates from MNNG/HOS orthotopic xenografts with or without Naa10p depletion were subjected to a Western blot analysis to determine MMP-2 expression levels. (**d**) An MMP-2-expressing plasmid (2 μg) was transfected into MNNG/HOS cells expressing shCtrl or shNaa10p as indicated and subjected to transwell invasion assays. Top: Expressions of MMP-2 and Naa10p are shown. Bottom: Quantitative results of the invasion assay are shown. Data were analyzed using a one-way ANOVA with Tukey’s post-hoc tests with 95% confidence intervals; different letters represent different levels of significance. (**e**) MMP-2 expression levels of MNNG/HOS cells expressing shCtrl or shNaa10p were detected by Western blotting at different time points after 35 μM cycloheximide treatment. Quantitative results of MMP-2 protein levels are shown at the bottom. (**f**) MNNG/HOS cells expressing shCtrl or shNaa10p with or without 10 μM MG-132 treatment for 6 h were harvested, and MMP-2, Naa10p, and β-actin expressions were analyzed by Western blotting. (**g**) Left: Representative microphotographs of immunohistochemical staining of Naa10p and MMP-2 in matched specimens of osteosarcomas. Scatter plots are shown in the right panel. The correlation between Naa10p and MMP-2 expressions was determined by a linear regression analysis.

### Naa10p directly interacts with MMP-2 through its acetyltransferase domain

To further investigate the mechanism of Naa10p in regulating MMP-2 protein stability, we next tested whether Naa10p was associated with MMP-2 in osteosarcoma cells. As shown in Figure 5a, MMP-2 protein was present in the immunocomplex precipitated by a Naa10p-specific antibody, and vice versa, in MNNG/HOS cells. In contrast, MMP-9 was not associated with Naa10p in MNNG/HOS cells (Supplementary Figure 1b). Naa15p also failed to coimmunoprecipitate MMP-2 in MNNG/HOS cells (Supplementary Figure 2), implying a highly specific interaction between Naa10p and MMP-2. We also purified GST-tagged Naa10p and incubated it with recombinant MMP-2 to examine if there was a direct interaction between them. Results from the pull-down assay clearly demonstrated that recombinant MMP-2 was directly associated with GST-Naa10p (Figure 5b). Furthermore, localization of Naa10p and MMP-2 was co-distributed in the cytosol as evaluated by an immunofluorescence staining analysis (Figure 5c). Naa10p was reported to interact with Naa15p through its N-terminal 58 residues and interact with different target proteins by the acetyltransferase domain or C-terminal region[16, 27]. To characterize the mechanism by which Naa10p and MMP-2 interact, we incubated recombinant MMP-2 with a series of truncated GST-fused Naa10p proteins. The recombinant MMP-2 protein could be pulled-down by full-length Naa10p, N-terminal-deleted Naa10p (residues 59~235), and the acetyltransferase domain (residues 60~130), but not by C-terminal Naa10p (residues 131~235) (Figure 5e). These results indicated that the acetyltransferase domain of Naa10p is the major binding region for MMP-2 interactions.

**Figure 5.**
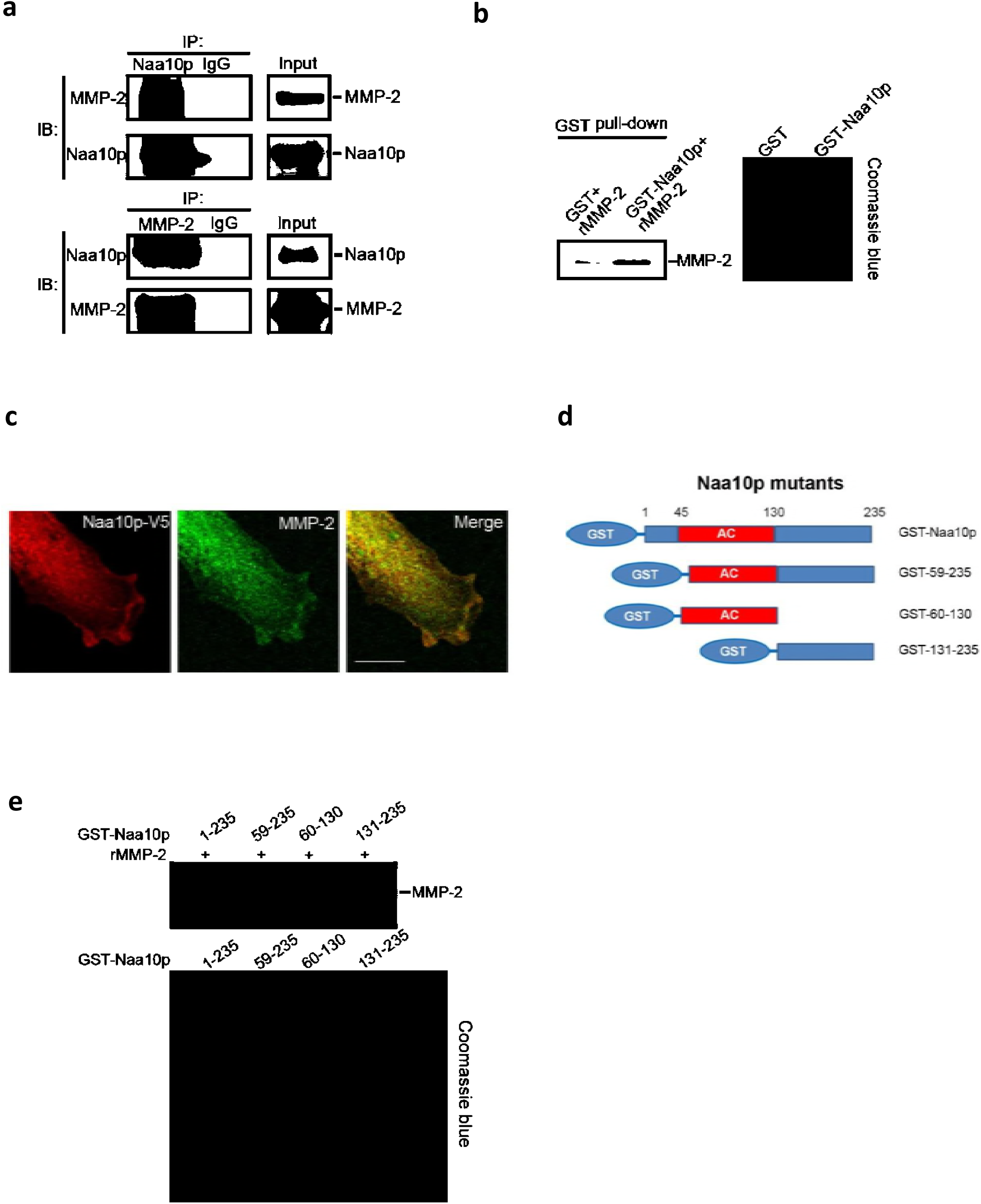
Naa10p directly interacts with matrix metalloproteinase (MMP)-2 through its acetyltransferase domain. (**a**) Western blot analysis of Naa10p and MMP-2 interactions in the immunocomplex precipitated from MNNG/HOS cell lysates with Naa10p (top) or MMP-2 (bottom) antibodies. Normal IgG antibodies were used as an IP control, and 10% whole-cell lysates were used as the input. (**b**) Western blot analysis of direct interaction of GST-Naa10p and recombinant MMP-2 protein after 16 h of co-incubation at 4 °C followed by pull-down with glutathione-Sepharose beads. Coomassie blue staining was used to show the loading amount. (**c**) Immunofluorescence staining of Naa10p and MMP-2 in MNNG/HOS cells with or without overexpressing Naa10p. Scale bar, 20 µm. (**d**) Schematic diagram demonstrates Naa10p constructs used in the GST-pull-down experiments. (**e**) Western blot analysis of the interaction regions of Naa10p and MMP-2 protein after GST-pull-down with different GST-Naa10p fragments and recombinant MMP-2. Coomassie blue staining was used to show the loading amount of GST-Naa10p fragments.

### N-terminal acetylation activity is essential for Naa10p-regulated MMP-2 stability and its cell invasive ability

Both acetylation- and non-acetylation-dependent regulation of Naa10p on its target proteins was previously reported.[12, 14, 16] To understand how Naa10p regulates MMP-2 protein stability, we compared the functional difference between wild-type Naa10p and the R82A mutant, an acetyl-CoA-binding site mutant which led to loss of enzyme activity in osteosarcoma cells. Unlike NAA10p, R82A overexpression in Saos-2 cells did not increase the MMP-2 protein level or invasive ability (Supplementary Figure 3). Furthermore, only re-expression of Naa10p, but not R82A in Naa10p-depleted MNNG/HOS cells significantly recovered the invasive ability of cells (Figure 6a). We also tested the rescue effect of Naa10p and R82A in an MNNG/HOS orthotopic model. Results showed that re-expression of Naa10p, but not R82A, also significantly restored lung metastasis and tumor growth abilities in MNNG/HOS/shNaa10p cell-injected mice (Figures 6b and Supplementary Figure 4). Since no detectable acetyl-lysine signal of MMP-2 was observed (data not shown), we focused on evaluating the effect of N-α-acetylation on the MMP-2 protein. The N-α-acetyltransferase activity of Naa10p required formation of the Naa10p-Naa15p complex, also known as the NatA complex. Knockdown of the NatA auxiliary subunit, Naa15p, also resulted in decreased MMP-2 expression and cell invasion in MNNG/HOS cells (Figure 6c). In addition, unlike overexpression of full-length Naa10p, overexpression of the N-terminal 58-residue-truncated Naa10p, which lacked Naa15p-binding capacity, did not induce an increase in MMP-2 expression (Figure 6d). Furthermore, we performed an N-terminal acetyltransferase assay on synthetic MMP-2 peptides with recombinant his-Naa10p and subjected them to mass spectrometry (MS) analysis. Under the same catalytic conditions, the methionine-removed MMP-2 peptide (EALMAR) showed seven-times more N-terminal acetylation than MMP-2 peptide (MEALMAR) (Supplementary Figure 5a). Tandem MS analysis confirmed α-acetylation on the second glutamate (Supplementary Figure 5b). Our results suggest that Naa10p may predominantly acetylate MMP-2 on the second glutamate. To verify the importance of N-terminal acetylation of MMP-2 in the maintenance of MMP-2 protein stability, we replaced the third residue of MMP-2 with proline (A3P-MMP-2) which was reported to prevent protein N-terminl-acetylation of other substrates[28, 29]. The protein half-life of A3P-MMP-2 in the presence of Naa10p-V5 overexpression, as determined by examining protein expression levels after CHX treatment, was shorter than that of wild-type MMP-2 (Figure 6e). Collectively, these data indicated that Naa10p forms a complex with Naa15p to acetylate MMP-2 in the N-terminus to maintain MMP-2 protein stability.

**Figure 6.**
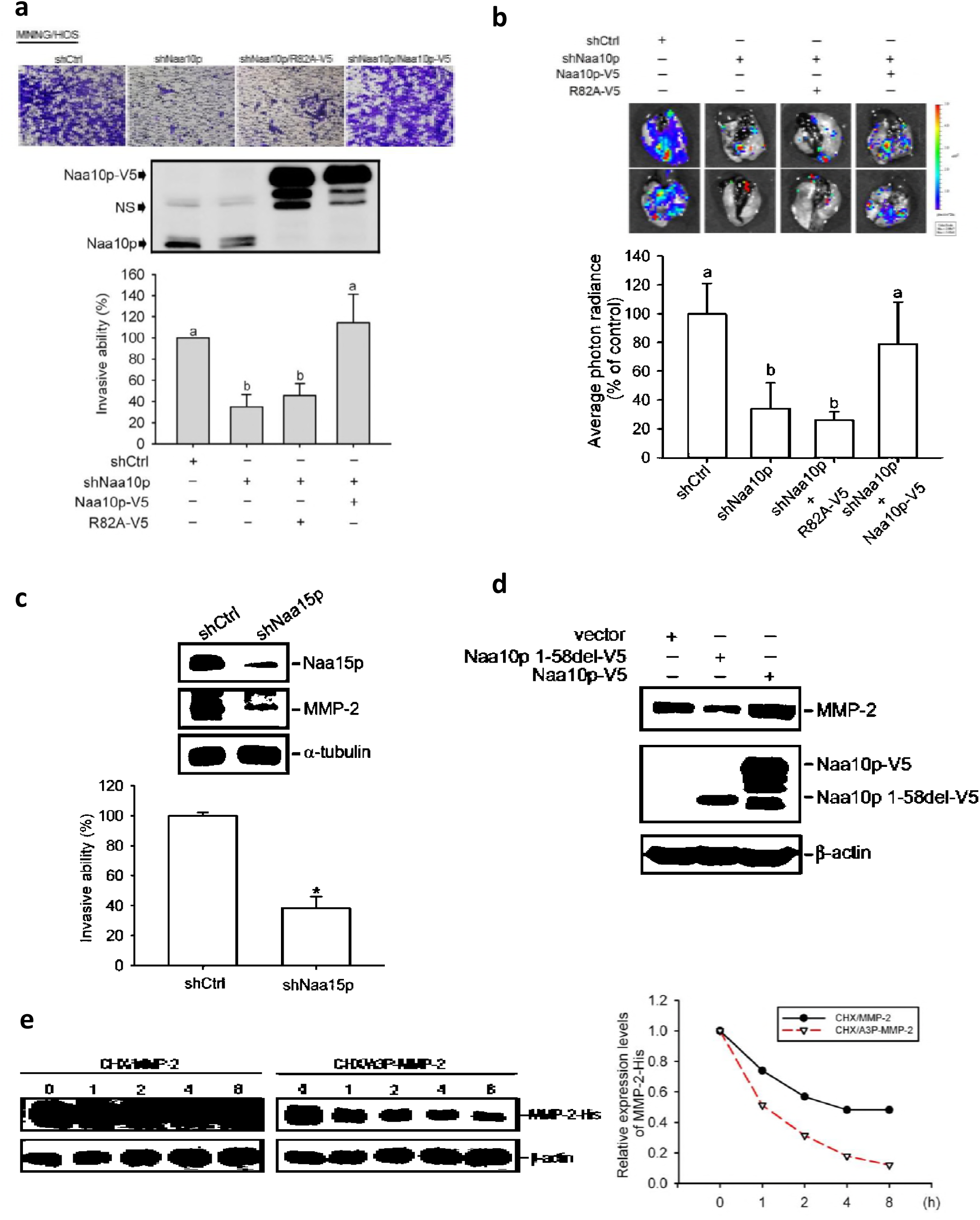
Naa10p regulates matrix metalloproteinase (MMP)-2 expression through N-terminal acetylation activity. (**a**) MNNG/HOS cells were infected with a lentivirus carrying shCtrl, shNaa10p, Naa10p-V5, or R82A-V5 as indicated. Top: Representative photos of transwell invasion assays. Middle: Western blot analysis of Naa10p expression levels. NS, non-specific bands. Bottom: Transwell invasion assays are shown as a percent of the control. (**b**) MNNG/HOS cells (5 × 10^5^) expressing shCtrl, shNaa10p, Naa10p-V5, or R82A-V5 as indicated were injected into the proximal tibia of NSG mice (six mice/group). Lung metastases were detected by measuring the bioluminescence activity. Bottom: Quantification of luciferase activity is shown as a percent of the shCtrl group. Data from (**a**) and (**b**) were analyzed using a one-way ANOVA with Tukey’s post-hoc tests with 95% confidence intervals; different letters represent different levels of significance. (**c**) MNNG/HOS cells were infected with a lentivirus carrying shCtrl or shNaa15p. Top: Western blot analysis of Naa15p and MMP-2 expression levels. Bottom: Results of the transwell invasion assays are shown as a percent of the control. Values are presented as the mean ± SE of three independent experiments. * *p* < 0.05, compared to the vehicle groups. (**d**) MNNG/HOS cells were transfected with a vector, Naa10p1-58del-V5, or Naa10p-V5 as indicated. Anti-MMP-2 and anti-V5 antibodies were used to detect MMP-2, Naa10p-V5, and Naa10p1-58del-V5 levels. (**e**) His-tagged MMP-2 or A3P-MMP-2 was transfected into MNNG/HOS cells. Left: His-tagged MMP-2 protein levels were monitored by Western blotting at different time points after 35 μM cycloheximide treatment. Right: Quantitative results of MMP-2 protein levels are shown.

## Discussion

MMPs are known for their role in promoting cancer progression and are considered ideal drug targets. However, MMPs’ intervention strategies have met with limited clinical success due to severe toxicities.[30] These poor results are likely due to the fact that MMPs play complex roles in both normal and cancer tissues. Most of the early programs targeting MMPs with broad-spectrum MMP-inhibitors were terminated due to poorly tolerated musculoskeletal pain and inflammation.[31] Therefore, for MMP-targeting to be clinically successful, they will need to be highly selective and able to accumulate in cancer tissues without eliciting adverse systemic effects. We identified that MMP-2 protein stability is tightly controlled by Naa10p, which is highly expressed in osteosarcoma tissues but rarely distributed in mature bone. Our findings revealed a possible targeting strategy by interfering with the Naa10p-MMP-2 regulatory axis in osteosarcomas.

Yoon et al. recently showed that Naa10p and Runx2 were simultaneously induced by bone morphogenetic protein (BMP)-2 during osteoblast differentiation.[22] BMP-2-induced Runx2 further stabilize the Naa10p protein through its inhibition of inhibitor of kappaB kinase (IKKβ)-mediated phosphorylation and degradation of Naa10p. It was also found that BMP-2 induces invasiveness of osteosarcoma mesenchymal stem cells by upregulating the extracellular matrix metalloproteinase inducer (EMMPRIN) and MMP-9.[32] Runx2 was reported to frequently be amplified and overexpressed in osteosarcomas[33] and promote invasion and metastasis of osteosarcoma cells in vitro[34, 35]. Recently, Runx-2 was also found to regulate MMP-2/9 expressions and cell invasion of osteosarcomas[36]. In this study, we identified that Naa10p was overexpressed in osteosarcoma cells, stabilized MMP-2 protein expression, and potentiated cell invasiveness and tumor metastasis. Although future evaluations are required, overexpression of Naa10p in osteosarcomas may be partially due to activation of BMP-2 and Runx-2 signaling. Moreover, whether Naa10p regulates MMP-2 expression and cell invasion by BMP-2 and Runx-2 also warrants further investigation.

Conflicting roles of Naa10p during cancer development were documented. For example, Kuo et al. reported that Naa10p acetylates the N-terminal methionine of TSC2, resulting in stabilization of the TSC1-TSC2 complex and suppression of the mammalian target of rapamycin (mTOR) pathway, eventually inhibiting cell proliferation and tumorigenesis.[12] Otherwise, a recent report identified Cdc25A as an Naa10p N-ε-acetylation target. The acetylated Cdc25A protein was stabilized and may thus accelerate G_1_/S and G_2_/M transitions, leading to genomic instability and promotion of tumorigenesis.[26] Consistently, Naa10p-transgenic mice were more sensitive to form oxidative tissue injury when exposed to hyperoxic conditions than their wild-type littermates, a phenomenon that may contribute to tumor initiation.[37] This evidence supports an oncogenic role for Naa10p in promoting tumorigenesis, although it is largely dependent on the affected targets. Although MMP-2 was rarely considered to affect tumor growth, we also observed a reduction in tumor volume in Naa10p-depleted osteosarcoma xenografts despite no significant difference in a short-term *in vitro* proliferation assay. Therefore, analysis of other Naa10p-target proteins in osteosarcomas may help reveal the molecular event by which Naa10p controls osteosarcoma growth *in vivo*.

The effects of Naa10p on cell invasiveness and metastasis are also complicated. In NSCLC, Naa10p was found to be expressed at a lower level in metastatic lymph nodes, compared to primary tumors. Accordingly, NSCLC patients with higher Naa10p expression also showed better prognoses with longer survival periods.[16] In contrast, Naa10p was reported to be a novel predictor of microvascular invasion in hepatocellular carcinoma.[38] Positive correlations of high Naa10p expression with lymph node metastasis and poor prognoses in breast cancer were also documented.[39] At the molecular level, Naa10p was found to acetylate myosin light chain kinase at lysine residues, and by doing so, deactivate MLCK, which led to decreased tumor cell migration and invasion.[13] Our previous study identified that the Naa10p-PIX association disrupted PIX-GIT-Paxillin complex formation in an acetylation-independent manner, and thus suppressed cell mobility.[16] The interaction targets of Naa10p are diverse and may highly depend on cancer types and cell contents. Furthermore, the distinct activities of Naa10p in regulating its associated proteins make its cellular functions even more diversified. Here in, we provide a novel mechanism to address the different face of Naa10p in regulating tumor metastasis. Our data suggest that MMP-2 is a novel N-terminal acetylation target of Naa10p. By acetylating and stabilizing the MMP-2 protein, Naa10p promotes osteosarcoma invasion and metastasis (Figure 7).

**Figure 7.**
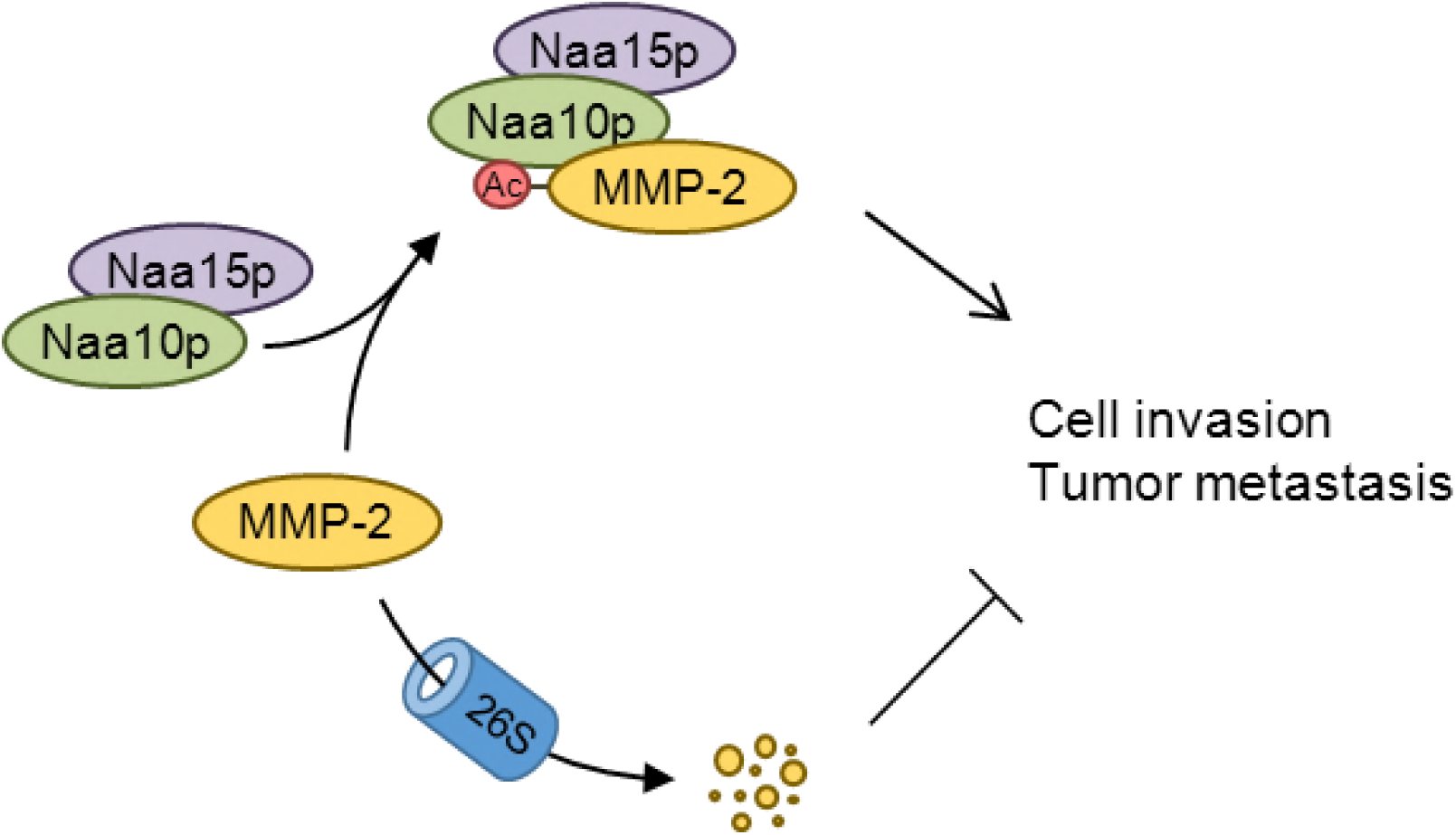
Schematic representation depicting the effects of Naa10p on matrix metalloproteinase (MMP)-2 stabilization. In invasive osteosarcoma cells, Naa10p was highly expressed and formed a complex with Naa15p to acetylate and stabilize the MMP-2 protein thus contributing to cell invasion and metastasis.

In conclusion, we showed that Naa10p is a poor prognostic factor the expression of which is correlated with metastasis and shorter survival in osteosarcomas. We propose that Naa10p can act as an oncogene by stabilizing MMP-2 protein expression in osteosarcomas. Our findings may provide a possible strategy for targeting Naa10p-MMP-2 regulation to prevent the highly metastatic property of osteosarcomas.

## Materials and Methods

### Materials

MG132 and cycloheximide (CHX) were purchased from Sigma-Aldrich (St Louis, MO). Cell culture materials and fetal bovine serum (FBS) were obtained from Gibco-BRL (Gaithersburg, MD). The following antibodies were used: Naa10p (GB-10511, Genesis Biotech, Taiwan) and MMP-2 (sc-13594, Santa Cruz Biotechnology, Santa Cruz, CA) for immunohistochemistry staining; Naa10p (GB-10511, Genesis Biotech) and MMP-2 (#4022, Cell Signaling Technology, Danver, MA, USA) for immunoblotting; Naa10p (sc-33820, Santa Cruz) and MMP-2 (sc-13594, Santa Cruz) for immunoprecipitation; Naa10p (GB-10511, Genesis Biotech) and MMP-2 (sc-13594 AF488, Santa Cruz) for immunofluorescence staining; and anti-V5 (#13202, Cell Signaling), anti-His (H1029, Sigma), and anti-Naa15p (A302-147A, Bethyl Laboratories, Montgomery, TX, USA). Naa10-V5 was a gift from Dr. Johan R. Lillehaug and Dr. Thomas Arnesen. The point mutation in the acetyltransferase domain of Naa10p (R82A) was made by site-directed mutagenesis using an overlapping polymerase chain reaction (PCR) followed by subcloning. Plasmids expressing the GST fusion proteins composed of full-length Naa10p, residues 59~235, 131~235, or 60~130 were engineered by a PCR, and subcloned into a pGEX-3X vector (GE Healthcare, Chicago, IL, USA). All DNA constructs were verified by DNA sequencing. The recombinant human MMP-2 was purchased from ProSpec-Tany TechnoGene (Rehovot, Israel). Unless otherwise specified, other chemicals used in this study were purchased from Sigma Chemical (St. Louis, MO).

### Cell culture

The human osteosarcoma cell lines, 143B, HOS, and MNNG/HOS, were purchased from American Type Culture Collection (authenticated using STR profile analysis, Manassas, VA, USA), while the U2OS, Saos-2, and MG-63 cell lines were obtained from the Food Industry Research and Development Institute (Hsinchu, Taiwan). 143B, HOS, and MNNG/HOS cells were maintained in α-minimum essential medium (MEM), and U2OS, Saos-2, and MG-63 cells were cultured in Dulbecco’s modified Eagle medium (DMEM). 10% fetal bovine serum (FBS), 2 mM L-glutamine, 100 U/mL penicillin, and 100 μg/mL streptomycin were added to both media and cells were cultured at 37 °C in a humidified atmosphere containing 5% CO_2_. All cell lines were confirmed to be Mycoplasma negative.

### *In vitro* wound-closure assay

MG-63 and Saos-2 cells (2 × 10^5^ cells/well) or HOS, MNNG/HOS, U2OS, and 143B cells (8 × 10^4^ cells/well) were seeded in 24-well plates for 24 h. After growth to confluence, the surface of the plate was scraped with a 200 -μL pipette tip to generate a cell-free zone, and then incubated with α-MEM or DMEM containing 10% FBS for 24 h. Cells were photographed using a phase-contrast microscope (100×) as previously described.[40]

### Western blot analysis

Protein lysates were prepared as described previously.[41] A Western blot analysis was performed with primary antibodies for Naa10p, Naa15p, MMP-2, V5, His, or β-actin. Image-Pro Plus software (Media Cybernetics, Silver Spring, MD) was used to quantify the density of the specific bands.

### Protease array analysis

Protein lysates from HOS/vector or HOS/shNaa10p cells were subjected to a protease (34 proteases) array analysis according to the manufacturer’s protocols (R&D Systems, Minneapolis, MN, USA). The pixel density of each spot on the developed x-ray film (Kodak, Perkin Elmer Life Sciences, Boston, MA) was analyzed using Image-Pro Plus software. Spot densities were normalized against respective reference array spots and then against controls.

### Cell viability assay

Osteosarcoma cell lines were stably transfected with either Naa10p-V5, Naa10p short hairpin (sh)RNA, or their respective controls and subjected to a cell viability assay (MTS assay; Promega, Madison, WI) according to the manufacturer’s instructions. Data were collected from three replicates.

### Transwell migration and invasion assays

We performed migration and invasion assays according to our previous study.[42] Briefly, 8 × 10^4^ and 10^5^ cells were respectively plated in an uncoated top chamber (24-well insert; pore size, 8 μm; Corning Costar, Corning, NY) for migration assay, and in a Matrigel (BD Biosciences, Bedford, MA)-coated top chamber for invasion assay. In both assays, cells were incubated in serum-free medium, and medium supplemented with serum was used for a chemoattractant in the lower chamber. After 24 h of incubation, the migrating cells on the lower surface of the membrane were fixed with methanol and stained with crystal violet. The number of cells invading through the membrane was counted under a light microscope (×100, three random fields per well).

### Lentiviral production and infection

The lentiviral vector and its packaging vectors were transfected into 293T packaging cells by calcium phosphate transfection. Briefly, 293T cells (10^6^) were split into 10-cm^2^ dishes 1 day before transfection. Then, cells were transfected with 10 μg pWPI-Naa10p or pWPI-control together with 10 μg of pCMVΔR8.91 (the packaging vector) and 1 μg of pMD.G (the envelope vector). After 5 h of incubation, the transfection medium was replaced with fresh culture medium. Forty-eight hours later, lentivirus-containing medium was collected from transfection and spun down at 1500 rpm for 5 min to pelletize the cell debris, the supernatant was filtered through a 0.45-um filter, and target cells were infected with fresh lentivirus-containing medium (supplemented with 8 μg/ml polybrene) for 24 h and subjected to different functional assays. Similarly, shNaa10 and shNaa15 were obtained from the National Core Facility for Manipulation of Gene Function by RNAi, miRNA, miRNA sponges, and CRISPR/Genomic Research Center, Academia Sinica. Lentiviral production and target cell infection were performed as described above.

### Construction of the Naa10-acetylated site-mutated MMP-2 plasmid and DNA transfection

The MMP-2 plasmid was obtained from GeneCopoeia (Rockville, MD). The point mutation in the third alanine residue of MMP-2 (A3P-MMP-2) was made by site-directed mutagenesis using an overlapping PCR followed by subcloning. To overexpress MMP-2 or A3P-MMP-2, semiconfluent cultures of HOS cells in a 6-cm Petri dish were transfected with 2 μg of an empty or expression vector, respectively, using Invitrogen Lipofectamine 2000 Transfection Reagent. After incubation for 24 h, cells were immunoblotted using an anti-MMP-2 or anti-His antibody to analyze the expression of exogenous MMP-2.

### Tissue array and immunohistochemistry (IHC)

The tissue arrays include an osteosarcoma and human normal bone tissue array (OS804, T261; BioMax, Rockville, MD). IHC staining for Naa10p and MMP-2 was performed on paraffin-embedded tissues. First of all, tissues were deparaffinized with xylene and incubated with 0.3% H_2_O_2_ to block endogenous peroxidase activity. Slides were washed with PBS and incubated with anti-Naa10p (1:200) or anti-MMP-2 (1:50) antibodies for 2 h at room temperature. Next, slides were thoroughly washed three times with PBS and were developed with a VECTASTAIN ABC (avidin-biotin complex) peroxidase kit (Vector Laboratories, Burlingame, CA) and a 3,3,9-diaminobenzidine (DAB) peroxidase substrate kit (Vector Laboratories) according to the manufacturer’s instructions. Nuclei were counterstained with hematoxylin. Naa10p and MMP-2 expression levels were semiquantitatively assessed in tissue samples as described previously. The tissues were scored in a blinded manner. A four-point staining intensity scoring system was devised to determine the relative expressions of Naa10p and MMP-2 in cancer specimens; the staining intensity score ranged from 0 (no expression) to 3 (maximal expression). Low expression was defined as no staining present (a staining intensity score of 0) or positive staining detected in < 20% of cells (a staining intensity score of 1); whereas high expression was defined as positive immunostaining present in 20%~50% of cells (a staining intensity score of 2) or >50% of the cells (a staining intensity score of 3).

### In vivo spontaneous metastasis model

All animal work was performed in accordance with protocols approved by the Institutional Animal Care and Use Committee of Taipei Medical University. Age-matched nonobese diabetic (NOD)-SCID male mice (6~8 weeks old) were used and randomly grouped for tumor growth and lung metastasis assays in an orthotopic graft model. 5 × 10^5^ luciferase-tagged MNNG/HOS cells expressing shNaa10p, Naa10p, R82A, or their respective controls were suspended in an 8:2 mixture of PBS and Matrigel, and implanted into the proximal tibia of mice (six mice per group). Luciferase-based, noninvasive bioluminescent imaging and analysis were performed with the Xenogen IVIS-200 system (Xenogen, Alameda, CA). Thirty-one days after tumor cell injection, animals were sacrificed, and tumor specimens were resected for a Western blot assay.

### In vivo experimental metastasis model

Age-matched NOD/SCID/IL2rγnull (NSG) male mice (6~8 weeks old) were used in the assays of tumor metastasis in an experimental metastasis model. Luciferase-tagged MNNG/HOS cells (10^6^) expressing shNaa10p or shCtrl were introduced into the circulation by injection into the tail vein with a 30-gauge needle. After 7 weeks, the difference in survival rates between these two groups (11 mice per group) was analyzed by the Kaplan-Meier test.

### In vitro acetylation assay

His-Naa10p was purified as previously described[43] and dissolved in 20mM Tris-HCl, 100mM NaCl, 1 mM EDTA, pH 8.0 solution. N-terminal peptides including MMP2 (MEALMAR) and de-MMP2 (EALMAR) were synthesized (Genomics, New Taipei city, TW). In the reaction mixture, 2µM enzyme, 500µM peptide substrate, and 1mM acetyl-CoA were added to the final 50μL solution. The *in vitro* acetylation mixture was incubated at 37°C for 25 minutes and stopped by adding stop solution (6M urea, 20mM Tris-HCl, 100mM NaCl, pH 8.0). After the *in vitro* acetylation reaction, the solutions were sent for the liquid chromatography–mass (LC-MS) spectrometry analysis for confirmation of the acetylated decoration on the peptides. High resolution LC-MS/MS was done by the LTQFT Ultra (Linear quadrupole ion trap Fourier transform ion cyclotron resonance) mass spectrometer (Thermo Electron, san Jose, CA) equipped with a nanoelectrospray ion source (New Objective, Inc.), an Agilent 1100 Series binary high-performance liquid chromatography pump (Agilent Technologies, Palo Alto, CA), and a FAMOS autosampler (LC Packings, San Francisco, CA). 6μl of reaction solution was injected at 10 μl/min flow rate onto a self-packed precolumn (150 µm I.D. × 20 mm, 5 µm, 100 Å). Chromatographic separation was performed on a self-packed reversed phase C18 nano-column (75 µm I.D. × 300 mm, 5 µm, 100 Å) using 0.1% formic acid in water as the mobile phase A and 0.1% formic acid in 80% acetonitrile as the mobile phase B, operated at a 300 nl/min flow rate. Survey full-scan MS conditions: mass range m/z 320-2000, resolution 100,000 at m/z 400.The ten most intense ions were sequentially isolated for MS2 by LTQ. Electrospray voltage was maintained at 1.8 kV and capillary temperature was set at 200 °C.

### Immunoprecipitation and GST-pull-down assays

MNNG/HOS cells were lysed in NETN buffer (20 mM Tris at pH 8.0, 100 mM NaCl, 1 mM EDTA, and 0.5% NP-40). Lysates (1 mg) were incubated with anti-Naa10p or anti-MMP-2 antibodies (5 μg) for 16 h at 4 °C followed by a 1-h incubation with 25 μl immobilized Protein A Sepharose beads. GST-Naa10p and truncated fragments were expressed and purified from *Escherichia coli* BL21. Recombinant MMP-2 (ENZ-769) was purchased from ProSpec (Ness-Ziona, Israel). GST-Naa10p or fragments (20 µg) and 2 µg rMMP2 were incubated in 1 ml NETN buffer for 16 h at 4 °C followed by pull-down with glutathione-Sepharose. Protein complexes collected from IP or pull-down were washed five times with 1 ml of NETN buffer, after which proteins were resolved on sodium dodecylsulfate polyacrylamide gel electrophoresis (SDS-PAGE) and analyzed by immunoblotting.

### Immunofluorescence staining

HOS/vector or HOS/Naa10p cells were grown on coverslips and fixed in 4% paraformaldehyde. Cells were permeabilized with 1% Triton X-100; labeled with a primary antibody against MMP-2 (1:100) (Alexa Fluor^®^ 488 conjugated), Naa10p (1:200), or V5 (1:200); and incubated with a secondary antibody conjugated to rhodamine (Santa Cruz Biotechnology, Santa Cruz, CA). Slides were examined and images captured using the Confocal Microscope and Single Molecule detection system (Leica, Germany).

### Statistical analysis

Data are presented as the mean ± SE. The statistical analysis was performed using Statistical Package for Social Science software, vers. 16 (SPSS, Chicago, IL). Student’s *t*-test was used to compare two groups. A one-way analysis of variance (ANOVA) followed by Tukey’s post-hoc test was used to analyze three or more groups. Statistical analyses of clinicopathological data were performed by a linear regression analysis. Survival curves were obtained using the Kaplan-Meier method. *p* values of < 0.05 were considered statistically significant.

## Acknowledgements

This research was supported by grants from the Ministry of Science and Technology, Taiwan (104-0210-01-09-02, 105-0210-01-13-01, and 106-0210-01-15-02 to M. Hsiao, 105-2320-B-002-035-MY3 to K.T. Hua and 105-2320-B-038 -058 -MY3 to M.H. Chien). We also thank GRC Mass Core Facility of Genomics Research Center, Academia Sinica, Taipei, Taiwan for Mass spectrometry analyses.

## Authors’ Contributions

Conception and design: M.-H Chien, K.-T. Hua Development of methodology: W.-J. Lee, Y.-C. Yang, P. Tan, K.-F. Pan Acquisition of data (provided animals, acquired and managed patients, provided facilities, etc.): W.-J. Lee, Y.-C. Yang, P. Tan, K.-F. Pan, H.-C. Tsai, Analysis and interpretation of data (e.g., statistical analysis, biostatistics, computational analysis): H.-C. Tsai, C.-H. Hsu, M. Hsiao Writing, review, and/or revision of the manuscript: M.-H Chien, K.-T. Hua Administrative, technical, or material support (i.e., reporting or organizing data, constructing databases): M.-H Chien, C.-H. Hsu, M. Hsiao, K.-T. Hua Study supervision: M.-H Chien, K.-T. Hua

